# Genome stability is in the eye of the beholder: recent retrotransposon activity varies significantly across avian diversity

**DOI:** 10.1101/2021.04.13.439746

**Authors:** James D. Galbraith, R. Daniel Kortschak, Alexander Suh, David L. Adelson

## Abstract

Since the sequencing of the zebra finch genome it has become clear the avian genome, while largely stable in terms of chromosome number and gene synteny, is more dynamic at an intrachromosomal level. A multitude of intrachromosomal rearrangements and significant variation in transposable element content have been noted across the avian tree. Transposable elements (TEs) are a source of genome plasticity, because their high similarity enables chromosomal rearrangements through non-allelic homologous recombination, and they have potential for exaptation as regulatory and coding sequences. Previous studies have investigated the activity of the dominant TE in birds, CR1 retrotransposons, either focusing on their expansion within single orders, or comparing passerines to non-passerines. Here we comprehensively investigate and compare the activity of CR1 expansion across orders of birds, finding levels of CR1 activity vary significantly both between and with orders. We describe high levels of TE expansion in genera which have speciated in the last 10 million years including kiwis, geese and Amazon parrots; low levels of TE expansion in songbirds across their diversification, and near inactivity of TEs in the cassowary and emu for millions of years. CR1s have remained active over long periods of time across most orders of neognaths, with activity at any one time dominated by one or two families of CR1s. Our findings of higher TE activity in species-rich clades and dominant families of TEs within lineages mirror past findings in mammals.

**Author Summary:** Transposable elements (TEs) are mobile, self replicating DNA sequences within a species’ genome, and are ubiquitous sources of mutation. The dominant group of TEs within birds are chicken repeat 1 (CR1) retrotransposons, making up 7-10% of the typical avian genome. Because past research has examined the recent inactivity of CR1s within model birds such as the chicken and the zebra finch, this has fostered an erroneous view that all birds have low or no TE activity on recent timescales. Our analysis of numerous high quality avian genomes across multiple orders identified both similarities and significant differences in how CR1s expanded. Our results challenge the established view that TEs in birds are largely inactive and instead suggest that their variation in recent activity may contribute to lineage-specific changes in genome structure. Many of the patterns we identify in birds have previously been seen in mammals, highlighting parallels between the evolution of birds and mammals.

## Introduction

Following rapid radiation during the Cretaceous-Paleogene transition, birds have diversified to be the most species-rich lineage of extant amniotes (Jarvis et al. 2014; Ericson et al. 2006; Wiens 2015). Birds are of particular interest in comparative evolutionary biology because of the convergent evolution of traits seen in mammalian lineages, such as vocal learning in songbirds and parrots (Bradbury and Balsby 2016; Petkov and Jarvis 2012; Pfenning et al. 2014), and potential consciousness in corvids (Nieder et al. 2020). However in comparison to both mammals and non-avian reptiles, birds have much more compact genomes (Gregory et al. 2007). Within birds, smaller genome sizes correlate with higher metabolic rate and the size of flight muscles (Hughes and Hughes 1995; Wright et al. 2014). However, the decrease in avian genome size occurred in an ancestral dinosaur lineage over 200 Mya, well before the evolution of flight (Organ et al. 2007). A large factor in the smaller genome size of birds in comparison to other amniotes is a big reduction in repetitive content (Zhang et al. 2014).

The majority of transposable elements (TEs) in the chicken (*Gallus gallus*) genome are degraded copies of one superfamily of retrotransposons, chicken repeat 1 (CR1) (International Chicken Genome Sequencing Consortium 2004). The chicken has long been used as the model avian species, and typical avian genomes were believed to have been evolutionarily stable due to little variation in chromosome number and chromosomal painting showing little chromosomal rearrangement (Burt et al. 1999; Shetty et al. 1999). These initial, low resolution comparisons of genome features, combined with the degraded nature of CR1s in the chicken genome, led to the assumption of a stable avian genome both in terms of karyotype and synteny but also in terms of little recent repeat expansion (International Chicken Genome Sequencing Consortium 2004; Wicker et al. 2005). The subsequent sequencing of the zebra finch (*Taeniopygia guttata*) genome supported the concept of a stable avian genome with little repeat expansion, but revealed many intrachromosomal rearrangements and a significant expansion of endogenous retroviruses (ERVs), a group of long terminal repeat (LTR) retrotransposons, since divergence from the chicken (Warren et al. 2010; Ellegren 2010). The subsequent sequencing of 48 bird genomes by the Avian Phylogenomics Project confirmed CR1s as the dominant TE in all non-passerine birds, with an expansion of ERVs in oscine passerines following their divergence from suboscine passerines (Zhang et al. 2014). The TE content of most avian genomes has remained between 7-10% not because of a lack of expansion, but due to the loss and decay of repeats and intervening non-coding sequence through non-allelic homologous recombination, cancelling out genome size expansion that would have otherwise increased with TE expansion (Kapusta et al. 2017). Since then, hundreds of bird species have been sequenced, revealing variation in karyotypes, and both intrachromosomal and interchromosomal rearrangements (Hooper and Price 2017; Damas et al. 2018; Feng et al. 2020; Kretschmer et al. 2020a, 2020b). This massive increase in genome sequencing has similarly revealed TEs to be highly active in various lineages of birds. Within the last 10 million years ERVs have expanded in multiple lineages of songbirds, with the newly inserted retrotransposons acting as a source of structural variation (Suh et al. 2018; Boman et al. 2019; Weissensteiner et al. 2020). Recent CR1 expansion events have been noted in woodpeckers and hornbills, leading to strikingly more repetitive genomes than the “typical” 7-10%. Between 23% to 30% of woodpecker and hoopoe genomes are CR1s, however their genome size remains similar to that of other birds (Feng et al. 2020; Manthey et al. 2018; Zhang et al. 2014). While aforementioned research focusing on the chicken suggested CR1s have not recently been active in birds, research focusing on individual avian lineages has used both recent and ancient expansions of CR1 elements to resolve deep nodes in a wide range of orders including early bird phylogeny (Suh et al. 2011; Matzke et al. 2012; Suh et al. 2015), flamingos and grebes (Suh et al. 2012), landfowl (Kriegs et al. 2007; Kaiser et al. 2007), waterfowl (St John et al. 2005), penguins (Watanabe et al. 2006), ratites (Haddrath and Baker 2012; Baker et al. 2014; Cloutier et al. 2019) and perching birds (Treplin and Tiedemann 2007; Suh et al. 2017). These studies largely exclude terminal branches and, with the exception of a handful of CR1s in grebes (Suh et al. 2012) and geese (St John et al. 2005), the timing of very recent insertions across multiple species remains unaddressed.

An understanding of TE expansion and evolution is important as they generate genetic novelty by promoting recombination that leads to gene duplication and deletion, reshuffling of genes and major structural changes such as inversions and chromosomal translocations (Zhou and Mishra 2005; Bailey et al. 2003; Lim and Simmons 1994; Underwood and Choi 2019; Lee et al. 2008; Chuong et al. 2017). TEs also have the potential for exaptation as regulatory elements and both coding and noncoding sequences (Warren et al. 2015; Wang et al. 2017; Barth et al. 2020). *Ab initio* annotation of repeats is necessary to gain a true understanding of genomic repetitive content, especially in non-model species (Platt et al. 2016). Unfortunately, many papers describing avian genomes (Cornetti et al. 2015; Jaiswal et al. 2018; Laine et al. 2016) only carry out homology-based repeat annotation using the Repbase (Bao et al. 2015) library compiled from often distantly related model avian genomes (mainly chicken and zebra finch. This lack of *ab initio* annotation can lead to the erroneous conclusion that TEs are inactive in newly sequenced species (Platt et al. 2016). Expectations of low repeat expansion in birds inferred from two model species, along with a lack of comparative TE analysis between lineages is the large knowledge gap we addressed here. As CR1s are the dominant TE lineage in birds, we carried out comparative genomic analyses to investigate their diversity and temporal patterns of activity.

## Results

### Identifying potential CR1 expansion across birds

From all publicly available avian genomes, we selected 117 representative assemblies not under embargo and with a scaffold N50 above 20,000 bp (available at July 2019) for analysis (SI Table 1). To find all CR1s that may have recently expanded in the 117 genomes, we first used the CARP *ab initio* TE annotation tool. From the output of CARP, we manually identified and curated CR1s with the potential for recent expansion based on the presence of protein domains necessary for retrotransposition, homology to previously described CR1s, and the presence of a distinctive 3’ structure. To retrotranspose and hence expand, CR1s require endonuclease and reverse transcriptase domains within a single ORF, and a 3’ structure containing a hairpin and microsatellite which potentially acts as a recognition site for the reverse transcriptase (Suh et al. 2014; Suh 2015). If a CR1 identified from homology contained both protein domains and the distinctive 3’ structure, we classified it as a “full length” CR1. We next classified a full length CR1 as “intact” CR1 if the endonuclease and reverse transcriptase were within a single intact ORF. Using the full length CR1s and previously described avian and crocodilian CR1s in Repbase as queries (Green et al. 2014; International Chicken Genome Sequencing Consortium 2004; Warren et al. 2010), we performed iterative searches of the 117 genomes to identify divergent, low copy number CR1s which may not have been identified by *ab initio* annotation. We ensured the protein domains and 3’ structures were present throughout the iterative searches. Assemblies with lower scaffold N50s generally contained fewer full length CR1s and none in the lowest quartile contained intact CR1s (Figure 1). Outside of the lowest quartile, assembly quality appeared to have little impact on the proportion of intact, full length repeats. The correlation of the low assembly quality with little to no full length CR1s was seen both across all species and within orders.

**Figure 1:**
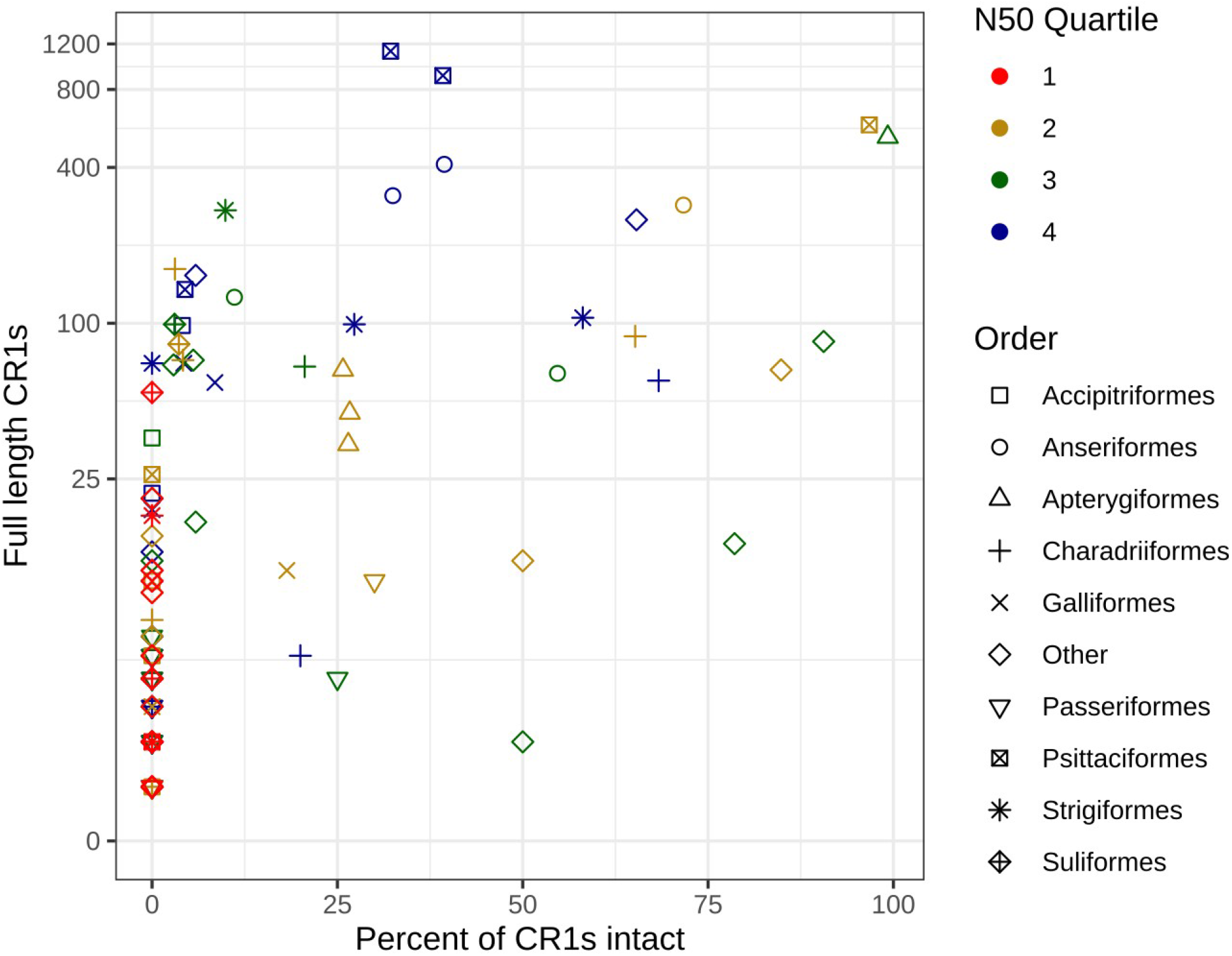
The impact of genome assembly quality on the identification of full length and intact CR1s. CR1s containing both an endonuclease and reverse transcriptase domains were considered full length, and those containing both domains within a single ORF considered intact. Both across all orders and within individual orders, genomes with higher scaffold N50 values (quartiles 2 through 4) had higher numbers of full length CR1s.

Our iterative search identified high numbers of intact CR1s in kiwis, parrots, owls, shorebirds and waterfowl (Figures 1 and 2). Only 2 of the 22 perching bird (Passeriformes) genomes contained intact CR1s, and all contained 10 or fewer full length CR1s. Similarly, of the 7 landfowl (Galliformes) genomes, only the chicken contained intact CR1s and contained fewer than 20 full length CR1s. High numbers of full length and intact repeats were also identified in two woodpeckers, Anna’s hummingbird, the chimney swift and the hoatzin, however, due to a lack of other genome sequences from their respective orders, we were unable to perform further comparative within order analyses of these species to look for recent TE expansion, i.e., within the last 10 million years. Of all the lineages we examined, only four have high quality assemblies of genera which have diverged within the last 10 million years and, based on the number of full length CR1s identified, the potential for very recent CR1 expansion: ducks (*Anas*), geese (*Anser*), Amazon parrots (*Amazona*) and kiwis (*Apteryx*) (Silva et al. 2017; Mitchell et al. 2014; Sun et al.2017). While the large number of full length repeats identified in owls is also high, we were unable to examine recent expansion in Strigiformes in detail due to the lack of a dated phylogeny. In addition to our genus scale analyses, we also examined CR1 expansion in parrots (Psittaciformes) overall, perching birds (Passeriformes) and shorebirds (Charadriiformes) since the divergence of each group, and compared the expansion in kiwis and their closest living relatives (Casuariiformes).

CR1 copy and those highlighted in yellow are the orders examined in detail. For coordinates of full length CR1s within genomes, see SI Data 1. Tree adapted from (Mitchell et al. 2014; Suh 2016).

**Figure 2:**
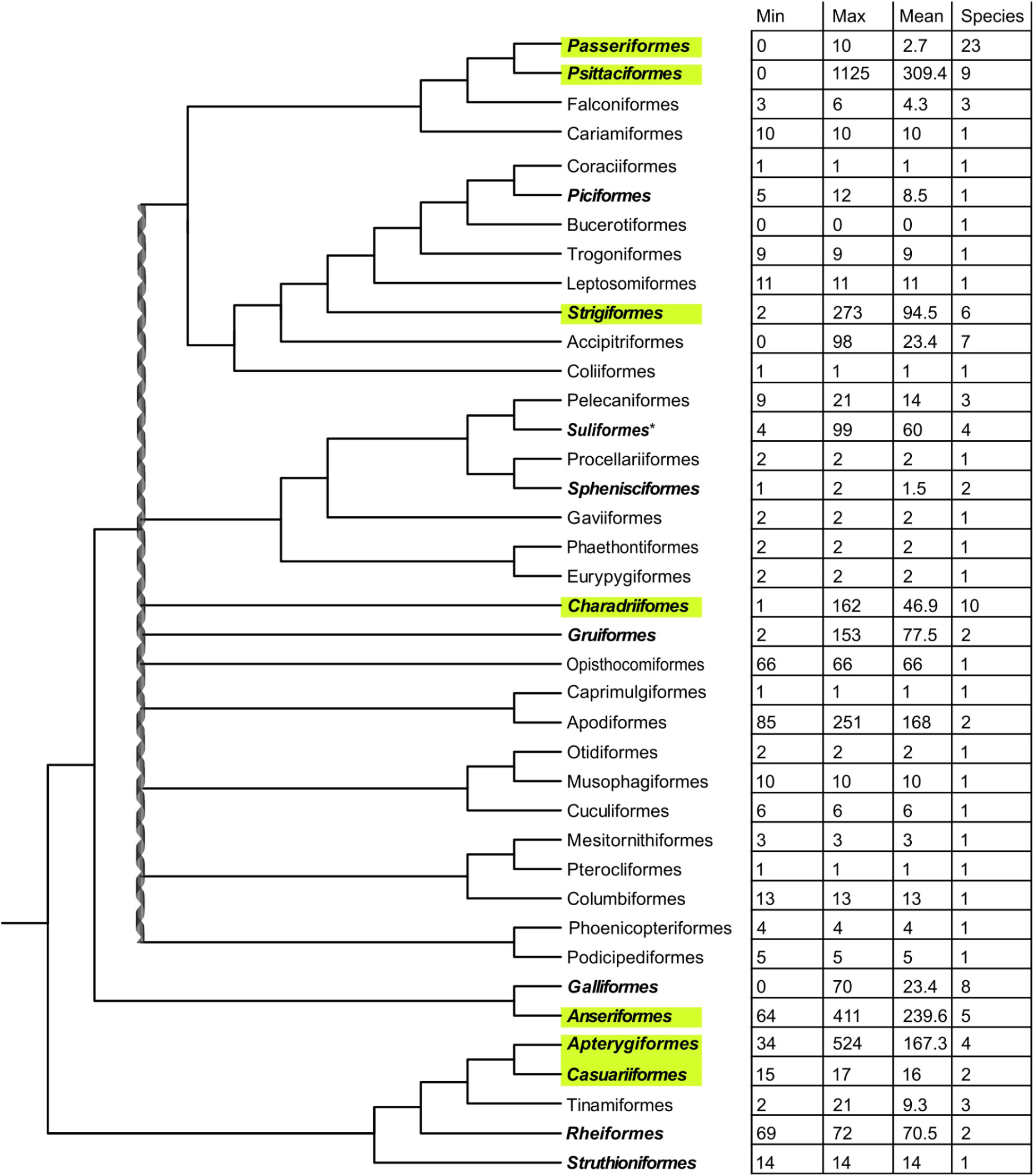
The number of full length CR1s varies significantly across the diversity of birds sampled. Minimum, maximum and mean number of full length CR1 copies identified in each order of birds, and the number of species surveyed in each order. Largest differences are noticeable between sister clades such as parrots (Psittaciformes) and perching birds (Passeriformes), and landfowl (Galliformes) and waterfowl (Anseriformes). The double helix represents a putative hard polytomy at the root of Neoaves (Suh 2016). Orders bolded contain at least one intact and potentially active

### Order-specific CR1 annotations and a phylogeny of avian CR1s reveal diversity of candidate active CR1s in neognaths

In order to perform comparative analyses of activity within orders, we created order-specific CR1 libraries. Instead of consensus sequences, all full length CR1s identified within an order were clustered and the centroids of the clusters were used as cluster representatives for that avian order. To classify the order-specific centroids, we constructed a CR1 phylogeny from the centroids and full length avian and crocodilian CR1s from Repbase (Figure 3, SI Figure 1, SI Data 2). From this tree, we partitioned CR1s into families to determine if groups of elements have been active in species concurrently. We partitioned the tree by eye based on the phylogenetic position of previously described CR1 families (Vandergon and Reitman 1994; Wicker et al. 2005; Warren et al. 2010; Bao et al. 2015) and long branch lengths rather than a cutoff for divergence, attempting to find the largest monophyletic groups containing as few previously defined CR1 families as possible. We took this “lumping” approach to our classification to avoid paraphyly and excessive splitting, resulting in some previously defined families being grouped together in one family (SI Table 2). For example, all full length CR1s identified in songbirds were highly similar to the previously described CR1-K and CR1-L families and were nested deeply within the larger CR1-J family. As a result, CR1-K, CR1-L and all full length songbird CR1s were reclassified as subfamilies of the larger CR1-J family. Based on the position of well resolved, deep nodes and previously described CR1s in the phylogeny, we defined 7 families of avian CR1s, with a new family, CR1-W, which was restricted to shorebirds. Interestingly, the 3’ microsatellite of the CR1-W family is a 10-mer rather than the octamer found in nearly all amniote CR1s (Suh 2015). With the exception of Palaeognathae (ratites and tinamous), all avian orders that contained large numbers of full length CR1s also contained full length CR1s from multiple CR1 families (Figure 3).

**Figure 3:**
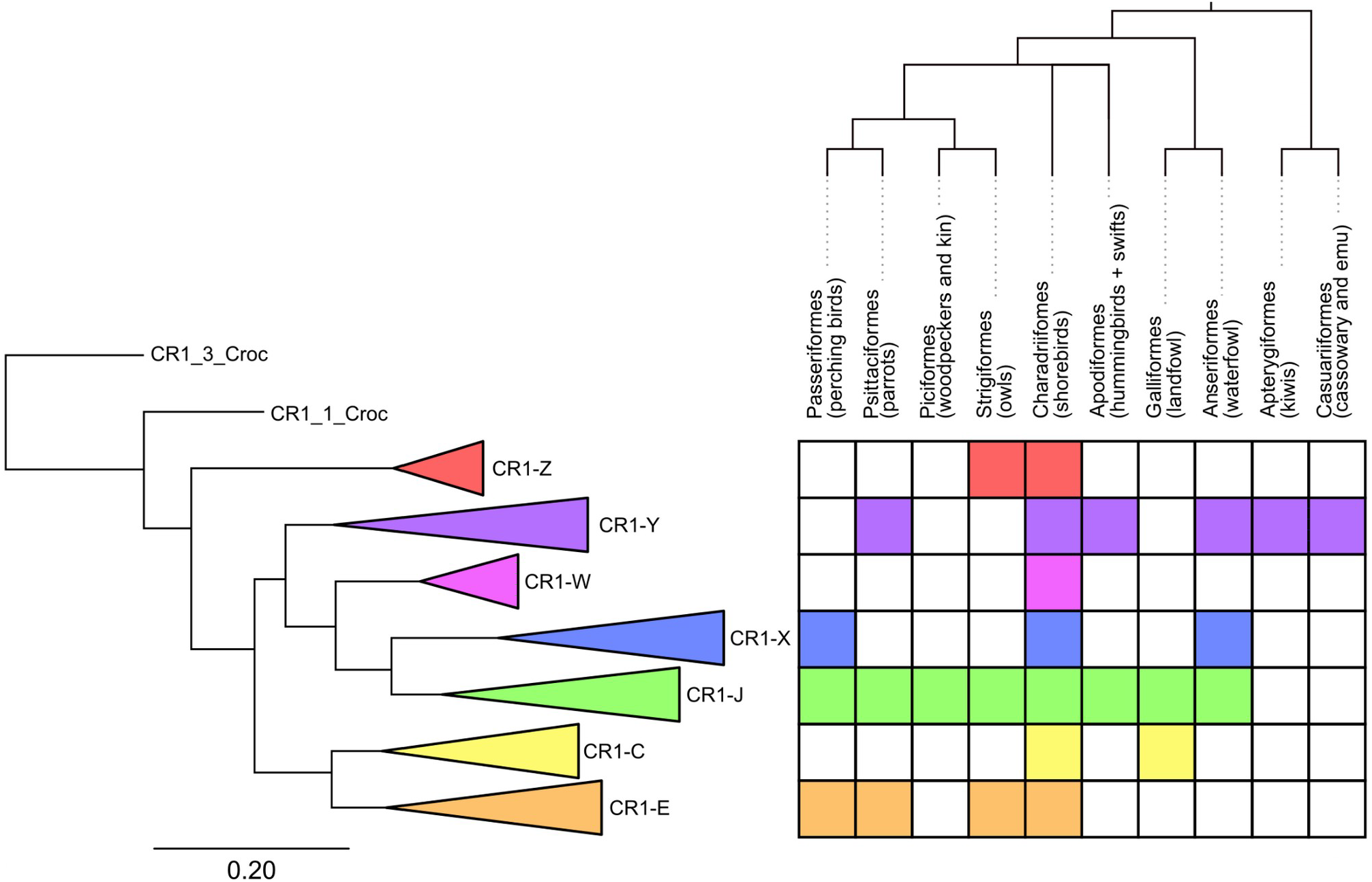
Collapsed tree of full length CR1s and presence of full length copies of CR1 families in selected avian orders. The name of each family is taken from a previously described CR1 present within the family (SI Table 3). The colouring of squares indicates the presence of full length CR1s within the order. All orders shown were chosen due to the presence of high numbers of intact CR1 elements, except for Casuariiformes which are shown due to their recent divergence from Apterygiformes as well as Passeriformes due to their species richness and frequent use as model species (especially zebra finch). The full CR1 tree was constructed using FastTree from a MAFFT alignment of the nucleotide sequences. For the full tree and nucleotide alignment of 1278 CR1s see SI Figure 1 and SI Data 2.

### Variable timing of expansion events across avian orders

We used the aforementioned order-specific centroid CR1s and avian and crocodilian Repbase sequences to create order-specific libraries. Throughout the following analysis we ensured CR1 copies identified were 3’ anchored, i.e. retain 3’ ends with homology to both the hairpin sequences and microsatellites. We used the order-specific libraries in reciprocal searches to identify and classify 3’ anchored CR1s present within all orders in which we had identified full length repeats. Using the classified CR1s we searched for all 3’ anchored CR1s (both full length and truncated) and constructed divergence plots to gain a basic understanding of CR1 expansions within each genome (SI Data 3). At high Jukes-Cantor distances, divergence profiles in each order show little difference between species. However, at lower Jukes-Cantor distances divergence, profiles differ significantly between species in some orders. For example, in songbirds at Jukes-Cantor distances higher than 0.1 the overall shape of the divergence plot curves and the proportions of the various CR1 families are nearly identical, while at distances lower than 0.1 higher numbers of the CR1-J family are present in some passerines than others (SI Figure 2a). CR1s most similar to all defined families were present in all orders of Galloanserae and Neoaves examined, with the exception of CR1-X which was restricted to Charadriiformes. Almost all CR1s identified in Palaeognathae genomes were most similar to CR1-Y with a small number of truncated and divergent repeats most similar to crocodilian CR1s (SI Data 3).

Divergence plots may not accurately indicate the timing of repeat insertions as they assume uniform substitution rates across the non-coding portion of the genome. High divergence could be a consequence of either full length CR1s being absent in a genome or the centroid identified by the clustering algorithm being distant from the CR1s present in a genome. To better determine when CR1 families expanded in avian genomes, we first identified regions orthologous to CR1 insertions sized 100-600 bp in related species (see Methods). We compared these orthologous regions and approximated the timing of insertion based on the presence or absence of the CR1 insertion in the other species. In most orders only long term trends could be estimated due to long branch lengths (cf. Figure 2) and high variability of the quality of genome assemblies (cf. Figure 1). Therefore, we focused our presence/absence analyses to reconstruct the timing of CR1 insertions in parrots, waterfowl, perching birds, and kiwis (Figure 4). We also applied the method to owls (SI Figure 3) and shorebirds (Figure 5), however due to the lack of order-specific fossil calibrated phylogenies of owls and long branch lengths of shorebirds, we could not determine how recent the CR1 expansions were.

**Figure 4:**
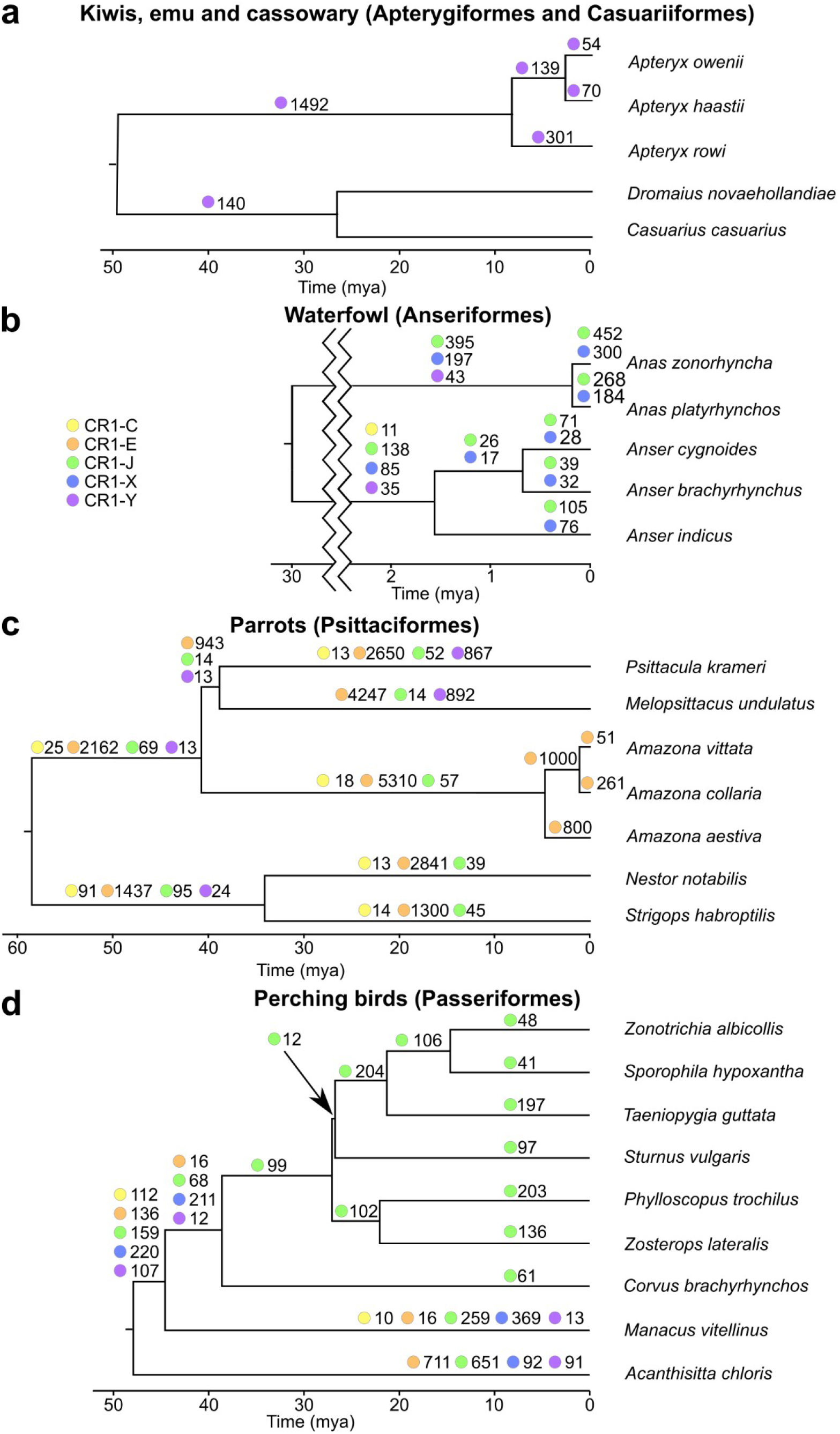
Presence/absence patterns reconstruct the timing of expansions of dominant CR1 families within five selected avian orders. The number next to the coloured circle is the number of CR1 insertions found. Only CR1 families with more than 10 CR1 presence/absence patterns (only CR1 insertions ranging between 100 and 600 bp were analyzed) are shown, for the complete number of insertions see SI Table 3. Phylogenies adapted from (Mitchell et al., 2014; Oliveros et al., 2019; Silva et al., 2017; Sun et al., 2017).

**Figure 5:**
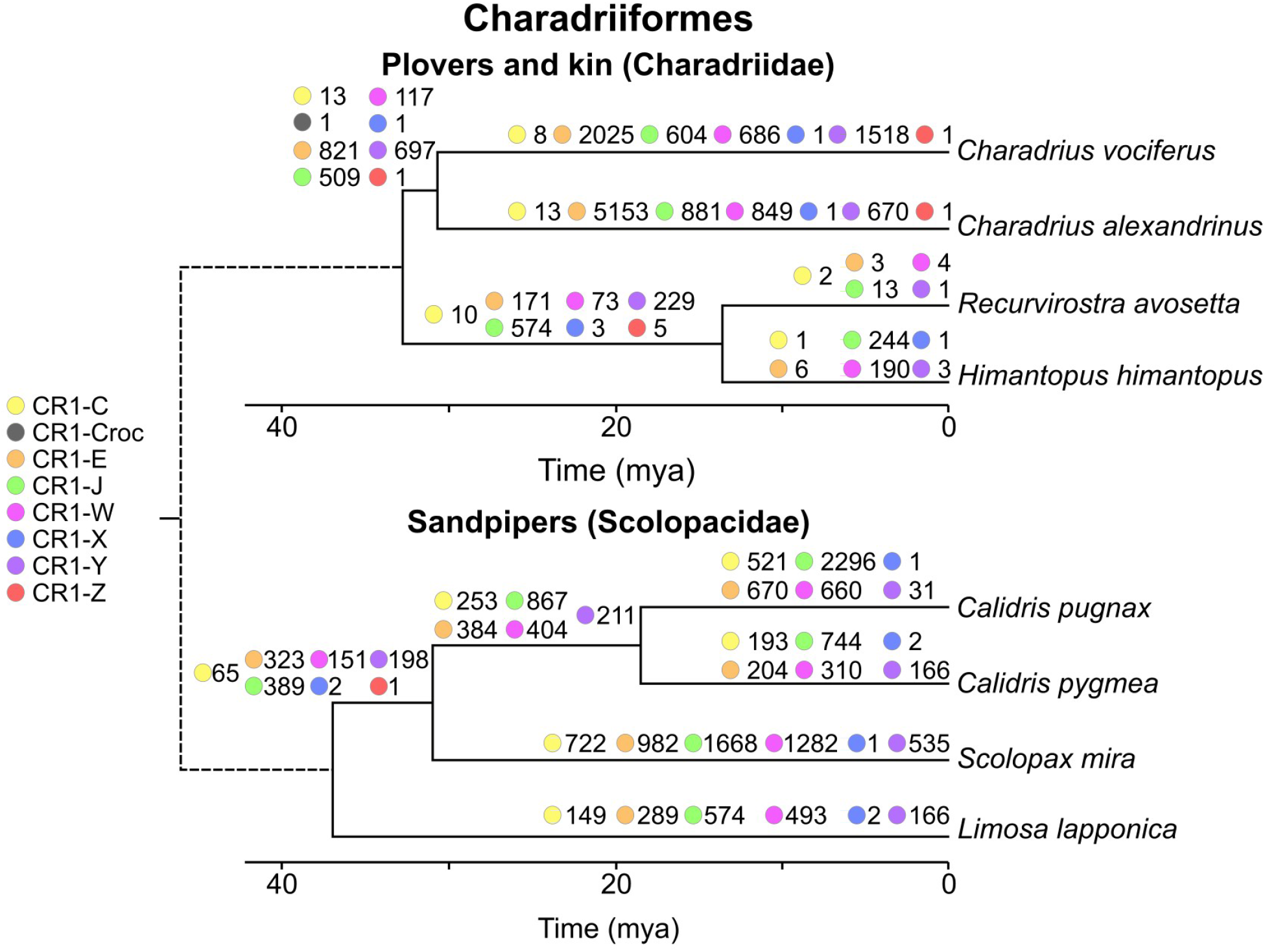
Presence/absence patterns reconstruct the timing of expansions of CR1 families in two lineages of shorebirds (Charadriiformes): plovers and sandpipers. The number next to the coloured circle is the number of CR1 insertions identified and only CR1 insertions between 100 and 600 bp long were analyzed. Divergence dates between plovers and sandpiper clades may differ due to the source phylogenies (Barth et al. 2013; Paton et al. 2003; Baker et al. 2007) being constructed using different approaches.

In analysing the repeat expansion in the kiwi genomes, we used the closest living relatives, the cassowary and emu (Casuariiformes), as outgroups. Following the divergence of kiwis from Casuariiformes, CR1-Y elements expanded, both before and during the recent speciation of kiwis over the last few My. In contrast, there was little CR1 expansion in Casuariiformes, both following their divergence from kiwis, and more recently since their divergence ∼28 Mya, with only 1 insertion found in the emu and 3 in the cassowary since they diverged (SI Table 3).

In the waterfowl species examined, both CR1-J and CR1-X families expanded greatly in both ducks and geese during the last 2 million years. Expansion occurred in both examined genera, with greater expansions in the ducks (*Anas*) than the geese (*Anser*). Other CR1 families appear to have been active following the two groups’ divergence ∼30 Mya, but have not been active since each genus speciated.

Due to the high number of genomes available for passerines, we chose best quality representative genomes from major groups *sensu (Oliveros et al. 2019)*; New Zealand wrens (*Acanthisitta chloris*), Suboscines (*Manacus vitellinus*), Corvides (*Corvus brachyrhynchos*), and Muscicapida (*Sturnus vulgaris*), Sylvida (*Phylloscopus trochilus* and *Zosterops lateralis*) and Passerida (*Taeniopygia guttata, Sporophila hypoxantha* and *Zonotrichia albicollis*). Between the divergence of Oscines (songbirds) and Suboscines from New Zealand wrens and the divergence of Oscines, there was a large spike in expansion of multiple families of CR1s, predominantly CR1-X. Since their divergence 30 Mya, only CR1-J remained active in oscines, though the degree of expansion varied between groups.

Of all avian orders examined, we found the highest levels of CR1 expansion in parrots. Because most branch lengths on the species tree were long, the timing of recent expansions could only be reconstructed in genus *Amazona*. The species from *Amazona* diverged 5 Mya ago and seem to vary significantly in their level of CR1 expansion. However, genome assembly quality might be a confounder as the number of insertions into a species of *Amazona* was highest in the best quality genome (*Amazona collaria*), and lowest in the worst quality genome (*Amazona vittata*). In all parrots, CR1-E was the predominant expanding CR1 family, however CR1-Y expanded in the *Melopsittacus-Psittacula* lineage, while remaining largely inactive in the other parrot lineages.

Multiple expansions of multiple families of CR1s have occurred in the two shorebird lineages examined; plovers (Charadriidae) and sandpipers (Scolopacidae) (Figure 5). The diversity of CR1 families that remained active through time was higher than in the other orders investigated, particularly in sandpipers, with four CR1 families showing significant expansion in *Calidris pugnax* and five in *Calidris pygmaea*, since their divergence. In all other orders examined in detail, CR1 expansions over similar time periods have been dominated by only one or two families, with insertions of fewer than 10 CR1s from non-dominant families (SI Table 2). Unfortunately, due to long branch lengths more precise timing of these expansions is not possible.

Finally, CR1s continuously expanded in true owls since divergence from barn owls, with almost all resolved insertions being CR1-E-like (SI Figure 3). However, due to the lack of a genus-level timed phylogeny, the precise timing of these expansions cannot be determined.

Combined, our CR1 presence/absence analyses demonstrate that the various CR1 families have expanded at different rates both within and across avian orders. These differences are considerable, ranging from an apparent absence of CR1 expansion in the emu and cassowary to slow, continued expansion of a single CR1 family in songbirds, to recent rapid expansions of one or two CR1 families in kiwis, Amazon parrots and waterfowl, as well as a wide variety of CR1 families expanding concurrently in sandpipers.

## Discussion

### Genome assembly quality impacts repeat identification

The quality of a genome assembly has a large impact on the number of CR1s identified within it, both full length and 5’-truncated. This is made clear when comparing the number of insertions identified within species in recently diverged genera. The three *Amazona* parrot species diverged approximately ∼2 Mya (Silva et al. 2017) and the scaffold N50s of *A. vittata, A. aestiva* and *A. collaria* are 0.18, 1.3 and 13 Mbp respectively. No full length CR1s were identified in *A. vittata*, and only 10 in *A. aestiva*, while 1125 were identified in *A. collaria*. Similarly, in *Amazona* the total number of truncated insertions identified increased significantly with higher scaffold N50s. In contrast the three species of kiwi compared, diverged ∼7 Mya and have similar N50s (between 1.3 and 1.7 Mbp). This pattern of higher quality genome assemblies leading to higher numbers of both full length and intact CR1s being identified is consistent across most orders examined, and is particularly true of the lowest N50 quartile (Figure 1). The lower number of repeats identified in lower quality assemblies is likely due to the sequencing technology used. Repeats are notoriously hard to assemble and are often collapsed, particularly when using short read Illumina sequencing, leading to fragmented assemblies (Alkan et al. 2011; Treangen and Salzberg 2011). The majority of the genomes we have used are of this data type. The recent sequencing of avian genomes using multiplatform approaches have resolved gaps present in short read assemblies, finding these gaps to be rich in interspersed, simple and tandem repeats (Peona et al. 2021; Li et al. 2021). Of particular note (Li et al. 2021) resolved gaps in the assembly of *Anas platyrhynchos* which we analyzed here using long read sequencing, and found the gaps to be dominated by the two CR1 families that have recently expanded in waterfowl (Anseriformes): CR1-J and CR1-X. Species with low quality assemblies may have full length repeats present in their genome, yet the sequencing technology used prevents the assembly of the repeats and hence detection. Thus TE activity may be even more widespread in birds than we estimate here.

### The origin and evolution of avian CR1s

Avian CR1s are monophyletic in regards to other major CR1 lineages found in amniotes (Suh et al. 2014). For comparison, crocodilians contain some CR1 families more similar to those found in testudines and squamates than others in crocodilians. By searching for truncated copies of previously described CR1s in addition to our order-specific CR1s, we were able to uncover how CR1s have evolved in avian genomes as birds have diverged. CR1-Y is the only family with full length CR1s present in Paleognathae, Galloanserae and Neoaves. The omnipresence of CR1-Y indicates it was present in the ancestor of all birds. A small number of highly divergent truncated copies of CR1s most similar to CR1-Z are found in ratites and CR1-J in tinamous (SI Figure 2b). This is potentially indicative of an ancestral presence of CR1-J and CR1-Z in the common ancestor of all birds, or misclassification owing to the high divergence of these CR1 fragments. As mentioned above, we took a lumping approach to classification to CR1 classification to avoid paraphyly, thereby collapsing highly similar families elsewhere considered as separate families. As CR1-C, CR1-E, and CR1-X are present in both Galloanserae and Neoaves but absent from Palaeognathae, we conclude these 4 families likely originated following the divergence of neognaths from paleognaths, but prior to the divergence of Neoaves and Galloanserae. In addition to having a 10 bp microsatellite instead of the typical 8 bp microsatellite, CR1-W is peculiar as it is unique to Charadriiformes but sister to CR1-J and CR1-X (Figure 3). This implies an origin in the neognath ancestor, followed by retention and activity in measurable numbers only in Charadriiformes.

A wide variety of CR1 families has expanded in all orders of neognaths, with many potential expansion events within the past 10 My present in many lineages. As mentioned in the results, it is not possible to conclude that insertions are ancient based on divergence plots alone. Some species with low quality genome assemblies, such as *A. vittata*, contained very few full length repeats compared to relatives (SI Figure 4). As a result of full length repeats not being assembled, the divergence of most or all truncated insertions identified in *A. vittata* would likely be calculated using CR1 centroids identified in *A. collaria*, leading to higher divergence values than those identified in *A. collaria*, and in turn an incorrect assumption of less recent expansion in *A. vittata* than *A. collaria*. In addition to fewer full length repeats being assembled, fewer truncated repeats also appear to have been assembled in poorer quality genomes.

### CR1 family expansions within orders

Across all sampled neognaths, recent expansions appear to be largely restricted to one or two families of CR1. Our presence/absence analyses found this to be the case in waterfowl, parrots, songbirds and owls, with shorebirds and the early passerine divergences the only exceptions. Similarly, based on the phylogeny of full length elements, most orders only retain full length CR1s from two or three families, while shorebirds retain full length CR1s from across all seven families. Our presence/absence analysis revealed likely concurrent expansions of at least four CR1 families in two families of shorebirds: sandpipers of genus *Calidris* and plovers of genus *Charadrius*. In both genera four families of CR1s have significantly expanded since their divergence including the order-specific CR1-W (Figure 5). While in both genera one family accounts for 40 to 50% of insertions, the other three families have hundreds of insertions each. This is highly different to the pattern seen in songbirds and waterfowl which, over a similar time period, have single digit insertions of non-dominant CR1 families (SI Table 3).

This increase of CR1 diversity in shorebirds could be due to some CR1 families in shorebirds having 3’ inverted repeat and microsatellite motifs which differ from the typical structure (Suh 2015) (SI Fig). For example, the CR1-W family has an extended 10 bp microsatellite (5’-AAATTCYGTG-3’) rather than the 8 bp microsatellite (5’-ATTCTRTG-3’) seen in nearly all other avian CR1s. When transcribed the 3’ structure upstream of the microsatellite is hypothesized to form a stable hairpin which acts as a recognition site for the cis-encoded reverse transcriptase (Suh 2015; Suh et al. 2017; Luan et al. 1993). The recently active CR1s we identified in other avian orders have 3’ microsatellites and hairpins which closely resemble those previously described. While the changes seen in shorebirds are minor we speculate they could impact CR1 mobilisation, allowing for more families to remain active than the typical one or two.

### Rates of CR1 expansion can vary significantly within orders

Based on the presence/absence of CR1 insertions and divergence plots, rates of CR1 expansion within lineages appear to vary even across rather short evolutionary timescales. The expansion of CR1-Y in kiwis appears to be a recent large burst of expansion and accumulation, while since Passeriformes diverged CR1-J appear to have continued to expand slowly in all families, however the number of new insertions seen in the American crow is much lower than that seen in the other oscine songbird species surveyed. The expansion of CR1-Y seen in the *Psittacula*-*Melopsittacus* lineage of parrots, following their divergence from the lineage leading to *Amazona*, appears to result from an increase in expansion, with little expansion in the period prior to divergence and none observed in other lineages of parrots. CR1s appear to have been highly active in all parrots examined since their divergence, however due to the less dense sampling it is not clear if this has been continuous expansion as in songbirds or a burst of activity like that in kiwis. Finally, in sandpipers CR1s have continued to expand in both species of *Calidris* since divergence, however the much lower number of new insertions in *C. pygmaea* suggests the rate of expansion differs significantly between the two species.

All full length CR1s identified in ratites were CR1-Y, and almost all truncated copies found in ratites were most similar to either CR1-Y, or crocodilian CR1s typically not found in birds (Suh et al. 2014). This retention of ancient CR1s and the presence of full length CR1s in species such as the southern cassowary (*Casuarius casuarius*) and emu (*Dromaius novaehollandiae*), yet without recent expansion, reflects the much lower substitution and deletion rates in ratites compared to Neoaves (Zhang et al. 2014; Kapusta et al. 2017). These crocodilian-like CR1s in ratites may be truncated copies of CR1s that were active in the common ancestor of crocodilians and birds (Suh et al. 2014) while we hypothesise that these have long since disappeared in Neoaves due to their higher deletion and substitution rates (Kapusta et al. 2017; Zhang et al. 2014).

### Co-occurrence of CR1 expansion with speciation

The four genera containing recent CR1 expansions we have examined co-occur with rapid speciation events. Of particular note, kiwis rapidly speciated into 5 distinct species composed of at least 16 distinct lineages arising due to significant population bottlenecks caused by Pleistocene glacial expansions (Weir et al. 2016). We speculate that the smaller population sizes might have allowed for CR1s to expand as a result of increased genetic drift (Szitenberg et al. 2016). While we do not see CR1 expansion occurring alongside speciation in passerines, ERVs, which are rare in other birds, have expanded throughout their diversification (Boman et al. 2019; Warren et al. 2010). Investigating the potentially ongoing expansion of CR1s and its relationship to speciation in ducks, geese, and Amazon parrots will require a larger number of genomes from within the same and sister genera to be sequenced, especially in waterfowl due to the high rates of hybridisation even between long diverged species (Ottenburghs et al. 2015).

### Comparison to mammals

As mentioned in the introduction, many parallels have been drawn between LINEs in birds and mammals, most notably the expansion of LINEs in both clades being balanced by a loss through purifying selection (Kapusta et al. 2017). Here we have found additional trends in birds previously noted in mammals. The TE expansion during periods of speciation seen in *Amazona, Apteryx* and *Anas* has previously been observed across mammals (Ricci et al. 2018). Similarly, the dominance of one or two CR1 families seen in most orders of birds resembles the activity of L1s in mammals (Ivancevic et al. 2016), however the general persistence of activity of individual CR1 families seems to be more diverse (Kriegs et al. 2007; Suh et al. 2011).

### Conclusion: the avian genome is more dynamic than meets the eye

While early comparisons of avian genomes were restricted to the chicken and zebra finch, where high level comparisons of synteny and karyotype led to the conclusion that bird genomes were largely stable compared to mammals (Ellegren 2010), the discovery of many intrachromosomal rearrangements across birds (Hooper and Price 2017; Skinner and Griffin 2012; Zhang et al. 2014; Farre et al. 2016) and interchromosomal recombination in falcons, parrots and sandpipers (O’Connor et al. 2018; Coelho et al. 2019; Pinheiro et al. 2021) has shown that at a finer resolution for comparison, the avian genome is rather dynamic. The highly variable rate of TE expansion we have observed across birds extends knowledge from avian orders with “unusual” repeat landscapes, i.e., Piciformes (Manthey et al. 2018) and Passeriformes (Warren et al. 2010), and provides further evidence that the genome evolution of bird orders and species within orders differs significantly, even though synteny is often conserved. In our comprehensive characterization of CR1 diversity across 117 bird genome assemblies, we have identified significant variation in CR1 expansion rates, both within genera such as *Calidris* and between closely related orders such as kiwis and the cassowary and emu. As the diversity and quality of avian genomes sequenced continues to grow and whole genome alignment methods improve (Feng et al. 2020; Rhie et al. 2020), further analysis of genome stability based on repeat expansions at the family and genus level will become possible. While the chicken and zebra finch are useful model species, models do not necessarily represent diversity of evolutionary trajectories in nature.

## Methods and Materials

### Identification and curation of potentially divergent CR1s

To identify potentially divergent CR1s we processed 117 bird genomes downloaded from Genbank (Benson et al. 2015) with CARP (Zeng et al. 2018); see SI Table for species names and assembly versions. We used RPSTBLASTN (Altschul et al. 1997) with the CDD library (Marchler-Bauer et al. 2017) to identify protein domains present in the consensus sequences from CARP. Consensuses which contained both an endonuclease and a reverse transcriptase domain were classified as potential CR1s. Using CENSOR (Kohany et al. 2006) we confirmed these sequences to be CR1s, removing others, more similar to different families of LINEs, such as AviRTEs, as necessary.

Confirmed CR1 CARP consensus sequences were manually curated through a “search, extend, align, trim” method as described in (Galbraith et al. 2020) to ensure that the 3’ hairpin and microsatellite were intact. Briefly, this curation method involves searching for sequences highly similar to the consensus with BLASTN 2.7.1+ (Zhang et al. 2000), extending the coordinates of the sequences found by flanks of 600 bp, aligning these sequences using MAFFT v7.453 (Katoh and Standley 2013) and trimming the discordant regions manually in Geneious Prime v2020.1. The final consensus sequences were generated in Geneious Prime from the trimmed multiple sequence alignments by majority rule.

### Identification of more divergent and low copy CR1s

To identify more divergent or low copy number CR1s which CARP may have failed to identify, we performed an iterative search of all 117 genomes. Beginning with a library of all avian CR1s in Repbase (Bao et al. 2015) (see SI Table 2 for CR1 names and species names) and manually curated CARP sequences we searched the genomes using BLASTN (-task dc-megablast - max_target_seqs <number of scaffolds in respective genome>), selecting those over 2700 bp and retaining 3’ hairpin and microsatellite sequences. Using RPSTBLASTN we then identified the full length CR1s (those containing both endonuclease (EN) and reverse transcriptase (RT) domains) and combined them with the previously generated consensus sequences. We clustered these combined sequences using VSEARCH 2.7.1 (Rognes et al. 2016) (--cluster_fast --id 0.9) and combined the cluster centroids with the Repbase CR1s to use as queries for the subsequent search iteration. This process was repeated until the number of CR1s identified did not increase compared to the previous round. From the output of the final round, order-specific clusters of CR1s were constructed and cluster centroids identified.

### Tree construction

To construct a tree of CR1s, the centroids of all order-specific CR1s were combined with all full length avian and two crocodilian CR1s from Repbase and globally aligned using MAFFT (--thread 12 --localpair). We used FastTree 2.1.11 with default nucleotide parameters (Price et al. 2010) to infer a maximum likelihood phylogenetic tree from this alignment, and rooted the tree using the crocodilian CR1s. The crocodilian CR1s were used as an outgroup as all avian CR1s are nested within crocodilian CR1s (Suh et al. 2015). This tree was split into different families of CR1 by eye based on the presence of long branches from high confidence nodes and the position of the previously described CR1 families from Repbase. To avoid excessive splitting and paraphyly of previously described families a lumping approach was taken resulting in some previously distinct families of CR1 from Repbase being treated as members of families they were nested within (SI Table 3).

### Identification and classification of CR1s within species

To identify, classify and quantify divergence of all 3’ anchored CR1s present within species, order-specific libraries were constructed from the order-specific clusters and the full length avian and crocodilian Repbase CR1s. 3’ anchored sequences CR1s were defined as CR1s retaining the 3’ hairpin and microsatellite sequences. Using these libraries as queries we identified 3’ anchored sequences CR1s present in assemblies using BLASTN. The identified CR1s were then classified using a reciprocal BLASTN search against the original query library.

### Determination of presence/absence in related species

To reconstruct the timing of CR1 expansions we selected the identified 3’ anchored CR1 copies of 100 and 600 bp length in a species of interest and at least 600 bp from the end of a contig, extending the coordinates of the sequences by 600 bp to include the flanking region and extracting the corresponding sequences. If the flanking regions contained more than 25% unresolved nucleotides (‘N’ nucleotides) they were discarded.

Using BLASTN we identified homologous regions in species belonging to the same order as the species being analysed, and through the following process of elimination identified the regions orthologous to CR1 insertions and their flanks in the related species. At each step of this process of elimination, if an initial query could not be satisfactorily resolved, we classified it as unscorable (unresolved) to reduce the chance of falsely classifying deletions or segmental duplications as new insertion events. First, we classified all hits containing the entire repeat and at least 150 bp of each flank as shared orthologous insertions. Following this, we discarded all hits with outer coordinates less than a set distance (150 bp) from the boundary of the flanks and CR1s to remove hits to paralogous CR1s insertions. This distance was chosen by testing the effect of a range of distances from 300 bp through to 50 bp in increments of 50 bp on a random selection of CR1s first identified in *Anser cygnoides* and *Corvus brachyrhynchos* and searched for in other species within the same order. Requiring outer coordinates to higher values resulted in higher numbers orthologous regions not being resolved, likely due to insertions or deletions within flanks since divergence. Allowing for boundaries of 50 or 100 bp resulted in many CR1s having multiple potential orthologous regions at 3’ flanks, many of were false hits, only showed homology to the target site duplication and additional copies of the 3’ microsatellite sequence. Thus 150 bp was chosen, as it was the shortest possible distance at which a portion of the flanking sequence was always present.

Based on the start and stop coordinates of the remaining hits, we determined the orientation the hit was in and discarded any queries without two hits in the same orientation. In addition, any queries with more than one hit to either strand was discarded. From the remaining data we determined the distance between the two flanks. If the two flanks were within 16 bp of each other in the sister species and the distance between the flanks was near the same length of the query CR1, the insertion was classified as having occurred since divergence. If the distance between the ends of the flanks in both the original species and sister species were similar, the insertion was classified as shared. For a pictorial description of this process including the parameters used, see SI Figure 5. This process was conducted for other species in the same order as the original species. Finally, we determined the timing of each CR1 insertion event by reconciling the presence/absence of each CR1 insertion across sampled species with the most parsimonious placement on the species tree (SI Figure 6).

## Supporting information

Phylogeny Data

Divergence Plots

Bird Genomes Used

Previous CR1 Classifications

Presence-Absence Results

## Acknowledgments

We thank Valentina Peona, Jesper Boman, Julie Blommaert and Alastair Ludington for comments on an earlier version of this manuscript.

## SI Information

### Figures

SI Figure 1. Phylogenetic tree of newly identified full length CR1s and full length avian CR1s from Repbase. The full length CR1s used are the centroids of order specific clusters constructed using VSEARCH at 90% identity. Phylogeny constructed using FastTree from a MAFFT alignment of the nucleotide sequences.

SI Figure 2. Scaled divergence of 3’ anchored CR1s identified in a) selected passerines and b) selected paleognaths. CR1s were initially identified using a reciprocal BLAST search based on libraries consisting of RepBase avian and crocodilian repeats and the centroids of full length sequences identified within the order clustered in VSEARCH.

SI Figure 3. Number of high confidence insertions of dominant CR1 families in owls approximated by presence/absence patterns of orthologous CR1 insertions between 100 and 600 bp in length. CR1 subfamilies are labeled by colour (see legend). Phylogeny adapted from (Salter et al. 2020).

SI Figure 4. Scaled divergence of 3’ anchored CR1s identified in species of Amazon parrot (*Amazona*). CR1s were initially identified using a reciprocal BLAST search based on a consisting of RepBase avian and crocodilian repeats and the centroids of full length sequences identified within parrots clustered in VSEARCH.

SI Figure 5. Presence/absence workflow. 3’ anchored CR1 insertions in a genome between 100 and 600 bp (1) were identified with BLASTN and had coordinates extended to include 600 bp of flanking sequence at both the 5’ and 3’ ends (2). The resulting 1300-1800 bp long sequences were searched for in a related genome using BLASTN. Hits containing the entire insertion and at least 150 bp of each flank were treated as ancestral insertions (3). Hits to insertion not containing any flanking region, with hits to the flanking sequence on differing strands or multiple hits to a single flanking sequence far from each other were treated as unresolvable and discarded. Insertions having at least 150 bp of each flank in close proximity and one flank containing at least 90 bp of the insertion were treated as ancestral insertions of which part was deleted in the species being searched (4). Sequences remaining were either flanks in close proximity or flanks plus a portion of the CR1 insertion. The distance between the flanks potentially containing part of the insertion was calculated in both species, qdist in the query species and sdist in the related species (5). If qdist was greater or equal to the length of the original CR1 insertion (olen) minus the length of 3x the 3’ microsatellite monomer and sdist was within the length of 2x the 3’ microsatellite monomer the insertion was treated as since divergence (6). If qdist was within the length of 2x the 3’ microsatellite monomer and the sdist was greater than 90 bp the insertion was treated as ancestral (7). Any insertions not fitting these criteria were treated as unresolvable and discarded. This strict process was calibrated through adjusting variables and viewing resulting pairwise alignments between regions identified as orthologous, using the presence of target site duplications in the query species and if part of the CR1 insertion was present in the related species to determine if insertions had truly occurred in an orthologous region, erring on the side of discarding new insertions over misclassifying partially deleted ancestral insertions as new insertions.

SI Figure 6. Presence/absence resolution - Example of the method we used to resolve the presence/absence, and hence insertion timing, of each CR1 in a species (species a), two related species (species b and c) and an outgroup (species d). The CR1 insertion in question is represented in green, the flanking regions in black and the branches labelled 1-3. The branch in bold italics is the branch on which the insertion occurred. If an CR1 was present in species a through c we considered the repeat to have been inserted at branch 1 (i), if in a and b at branch 2 (ii) and if in species a alone to be since the divergence from the immediate sister species and on branch 3 (iii). If present in all three species and the outgroup species examined we consider the repeat to be ancestral (iv). If a CR1 was absent from an immediate sister species but present in the more distant related species we considered this to be a result of deletion in the immediate sister species (v). Finally, if the orthologous region was present in a species or group of species but could not be resolved in the immediate sister species we considered the timing of insertion to be unresolvable (vi).

### Tables

SI Table 1. Genome assemblies used throughout this analysis. All genomes were downloaded from GenBank.

SI Table 2 - Reclassification of previously described full length avian CR1s based on their position within our CR1 phylogeny (SI Figure 1; same color coding).

SI Table 3. Resolution of presence or absence of orthologous CR1 insertions between 100 and 600 bp in related species in waterfowl, shorebirds, perching birds, parrots, owls, and kiwis + cassowary + emu genomes. Cells highlighted in yellow are the values used to construct Figures 4 and 5 and SI Figure 3.

## Data

SI Data 1 - Coordinates of full length CR1s identified in each genome in BED format. For the appropriate genome version see SI Table 1.

SI Data 2 - Multiple sequence alignment used to create the CR1 phylogeny (SI Figure 1) and Newick tree of said phylogeny.

SI Data 3. Divergence plots of 3’ anchored CR1s identified in each species of bird belonging to orders in which we detected full length CR1s. CR1s were identified using a reciprocal BLAST search based on libraries consisting of Repbase avian and crocodilian repeats and the centroids of full length sequences identified within the order clustered in VSEARCH. Jukes-Cantor distance was calculated from the reciprocal BLAST search output.

