## Supplementary figures and images for "Genome stability is in the eye of the beholder: recent retrotransposon activity varies significantly across avian diversity"

### Fig_1_Activity_Across_Species.png

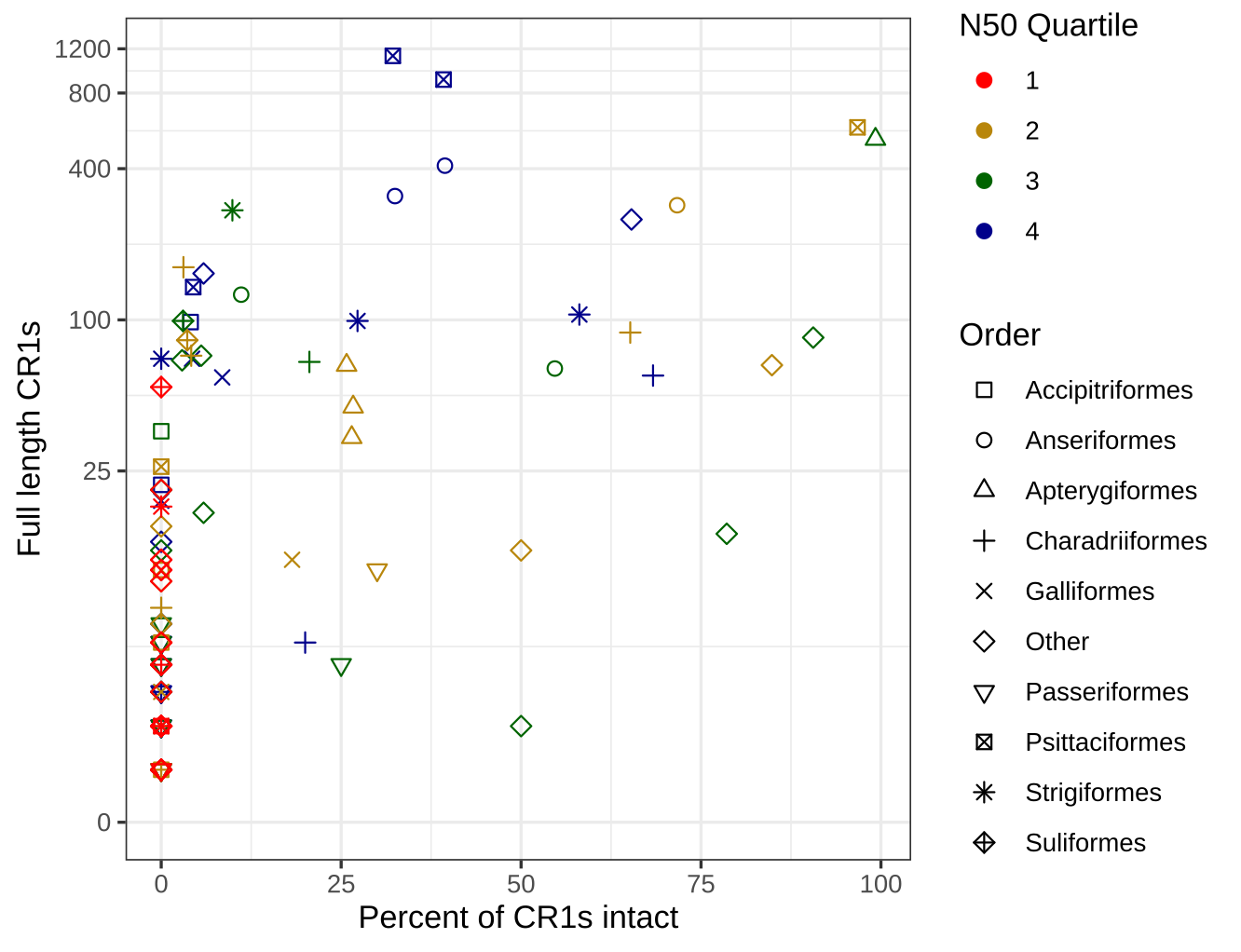

### Fig_2.png

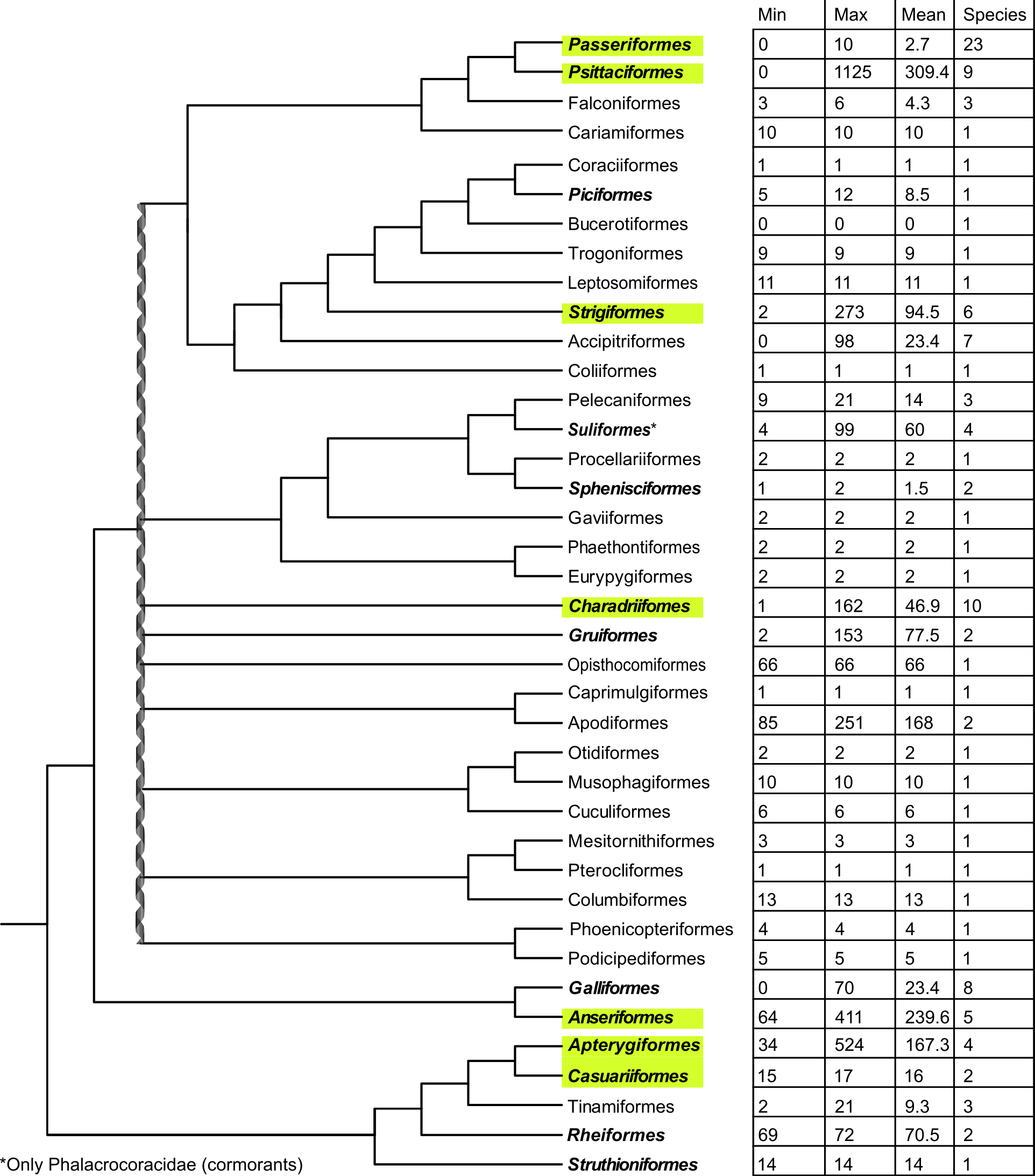

### Fig_3_RT_EN_90_collapsed_1.png

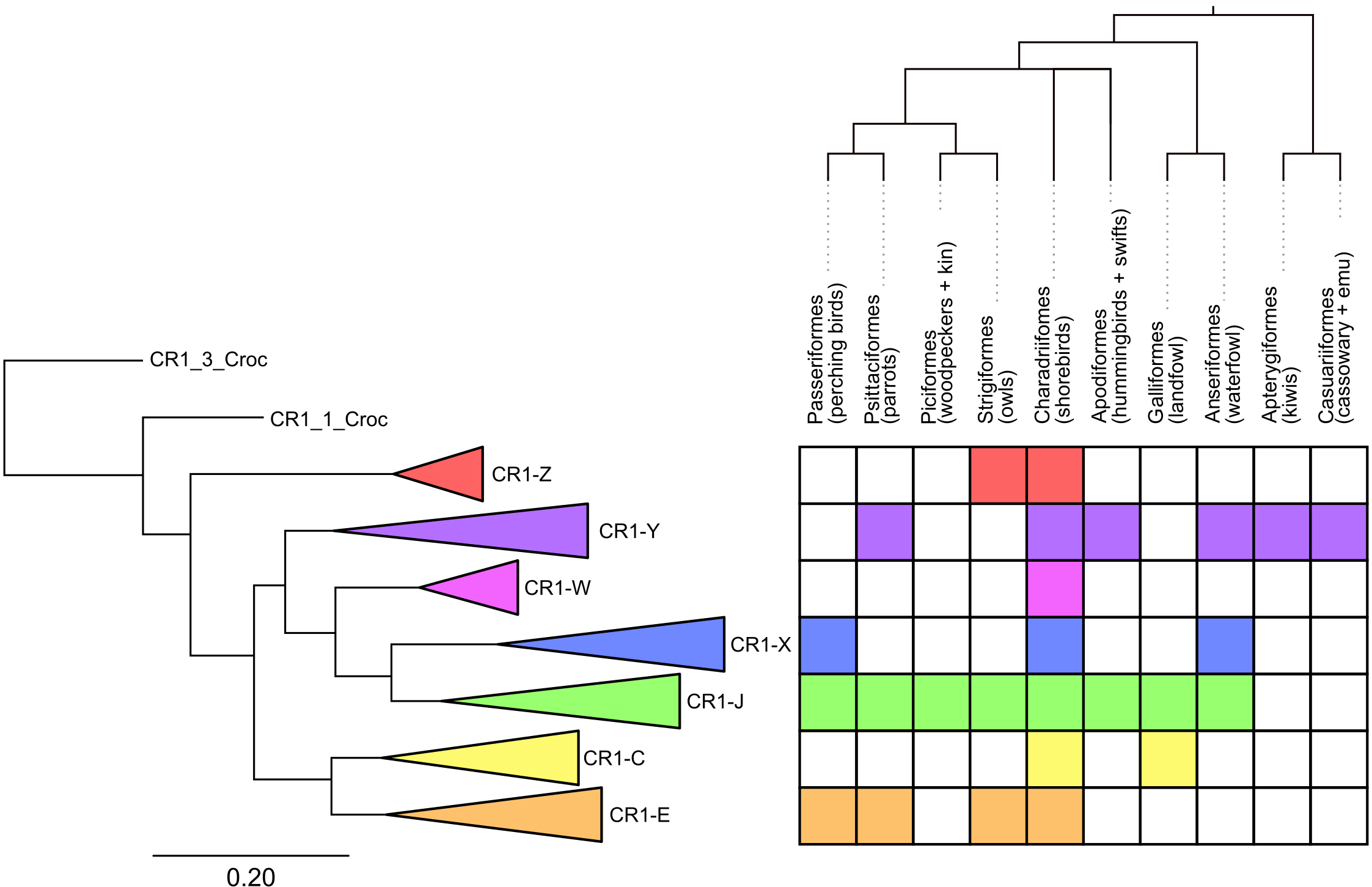

### Fig_4_compiled_insertions.png

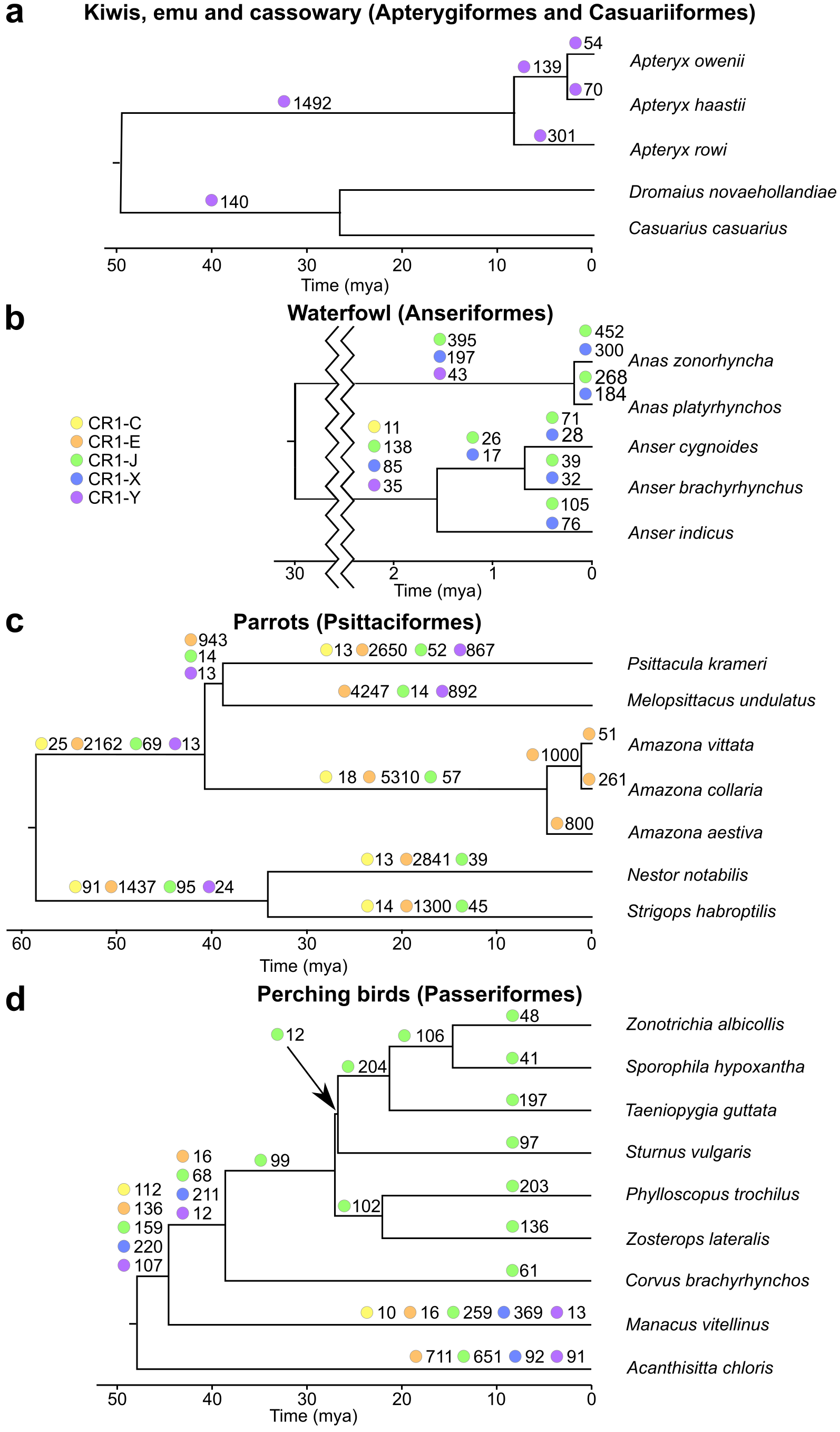

### Fig_5_shorebirds.png

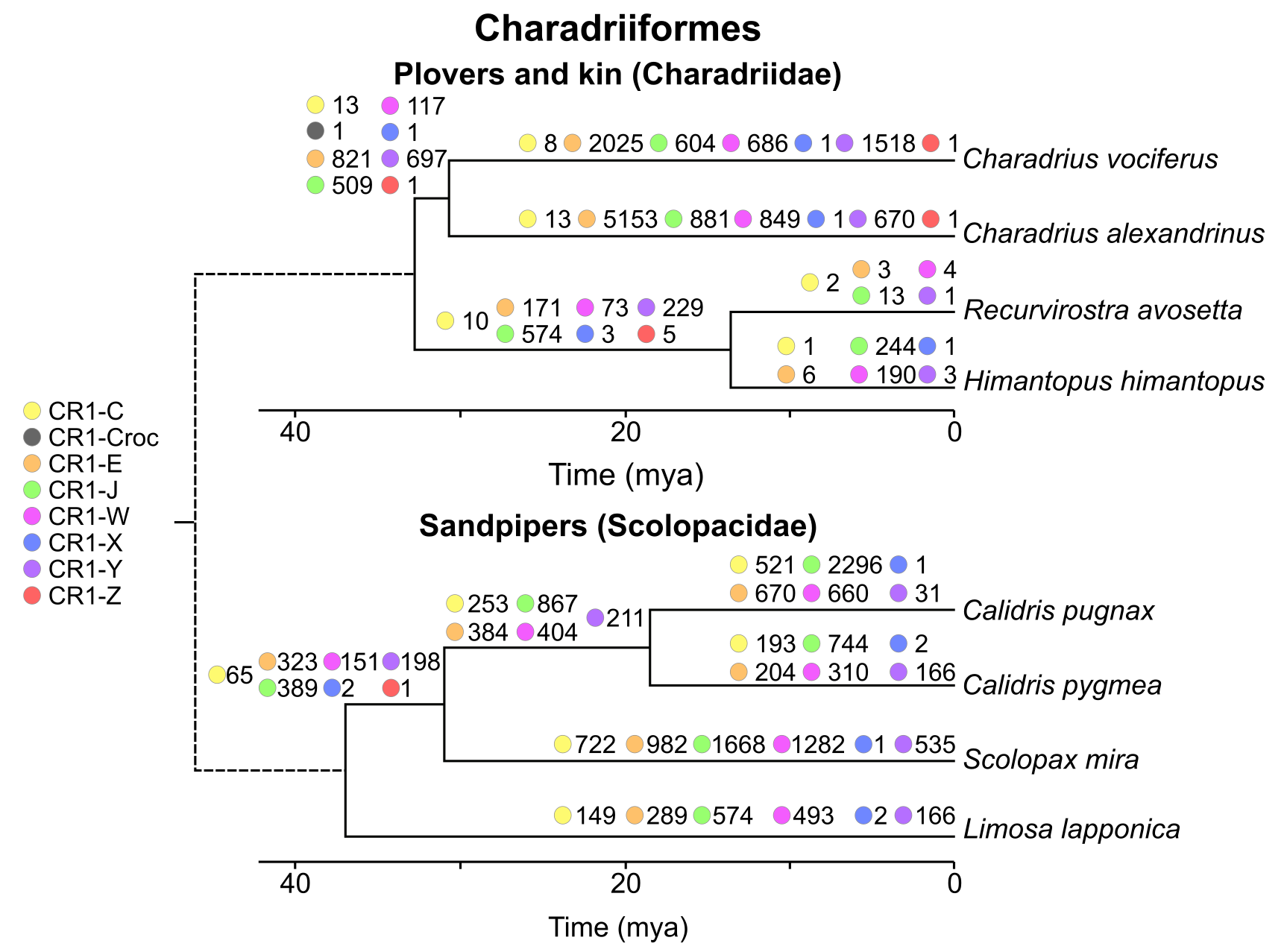

### SI_Fig_1_RT_EN_90_expanded.png

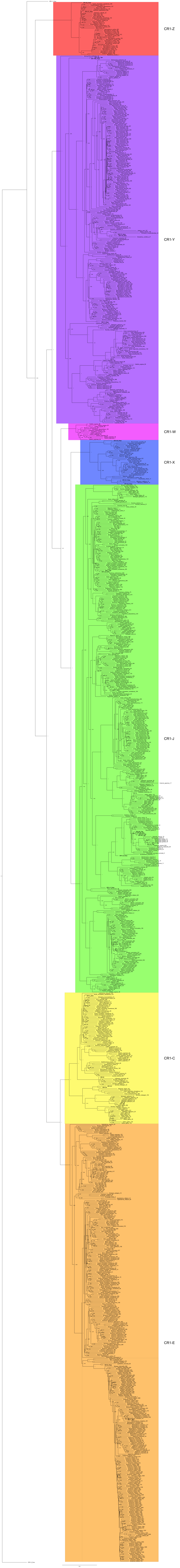

### SI_Fig_2_divergence_in_orders.png

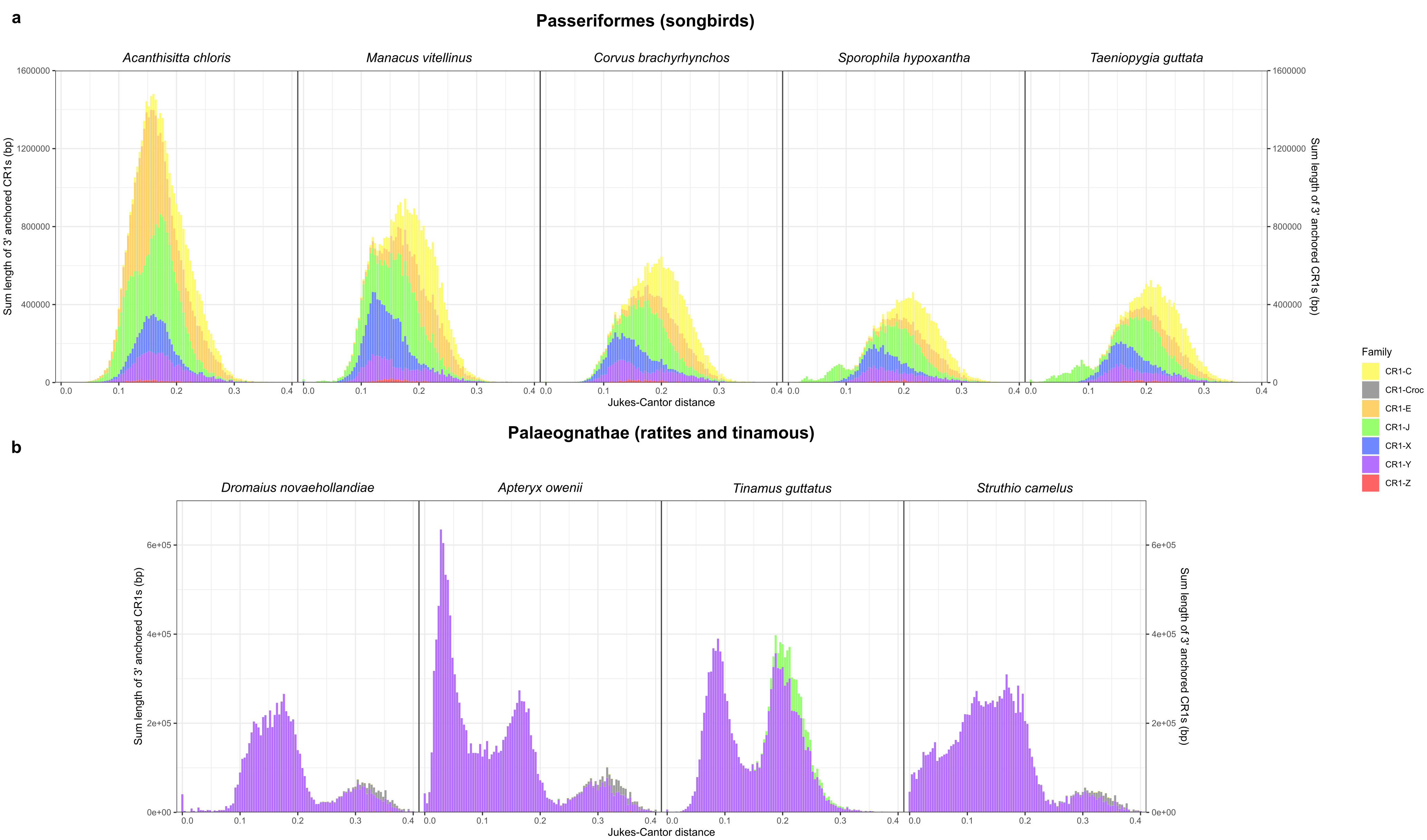

### SI_Fig_3_strigiformes.png

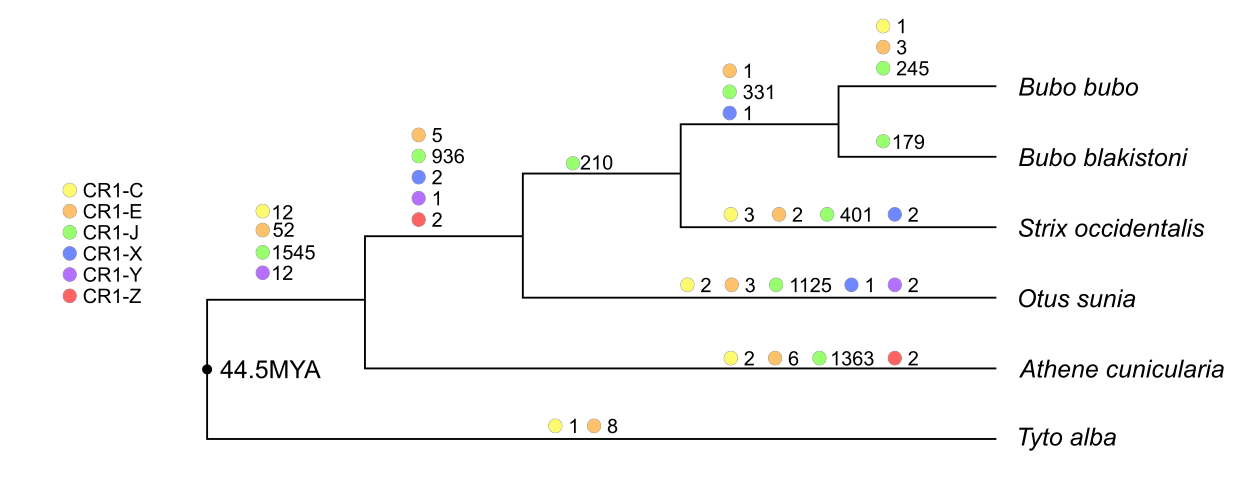

### SI_Fig_4_divergence_in_Amazona.png

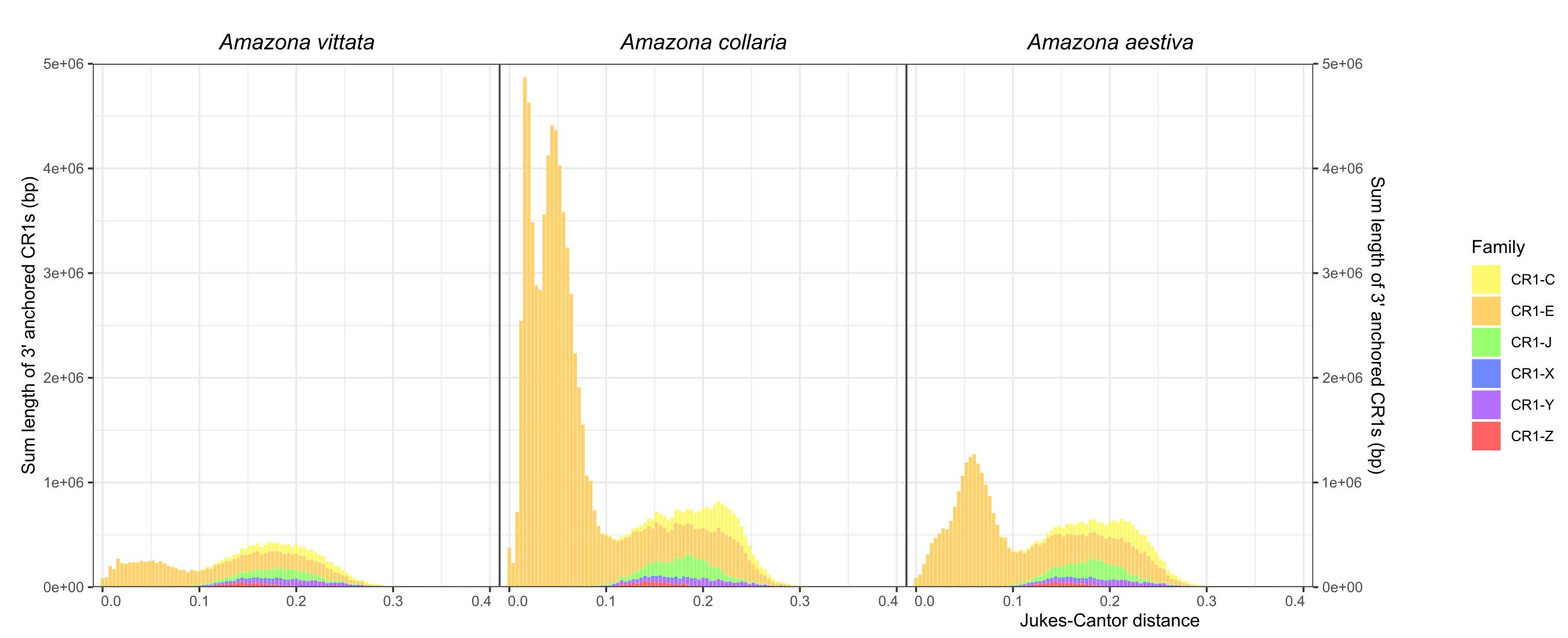

### SI_Fig_5_Technique.png

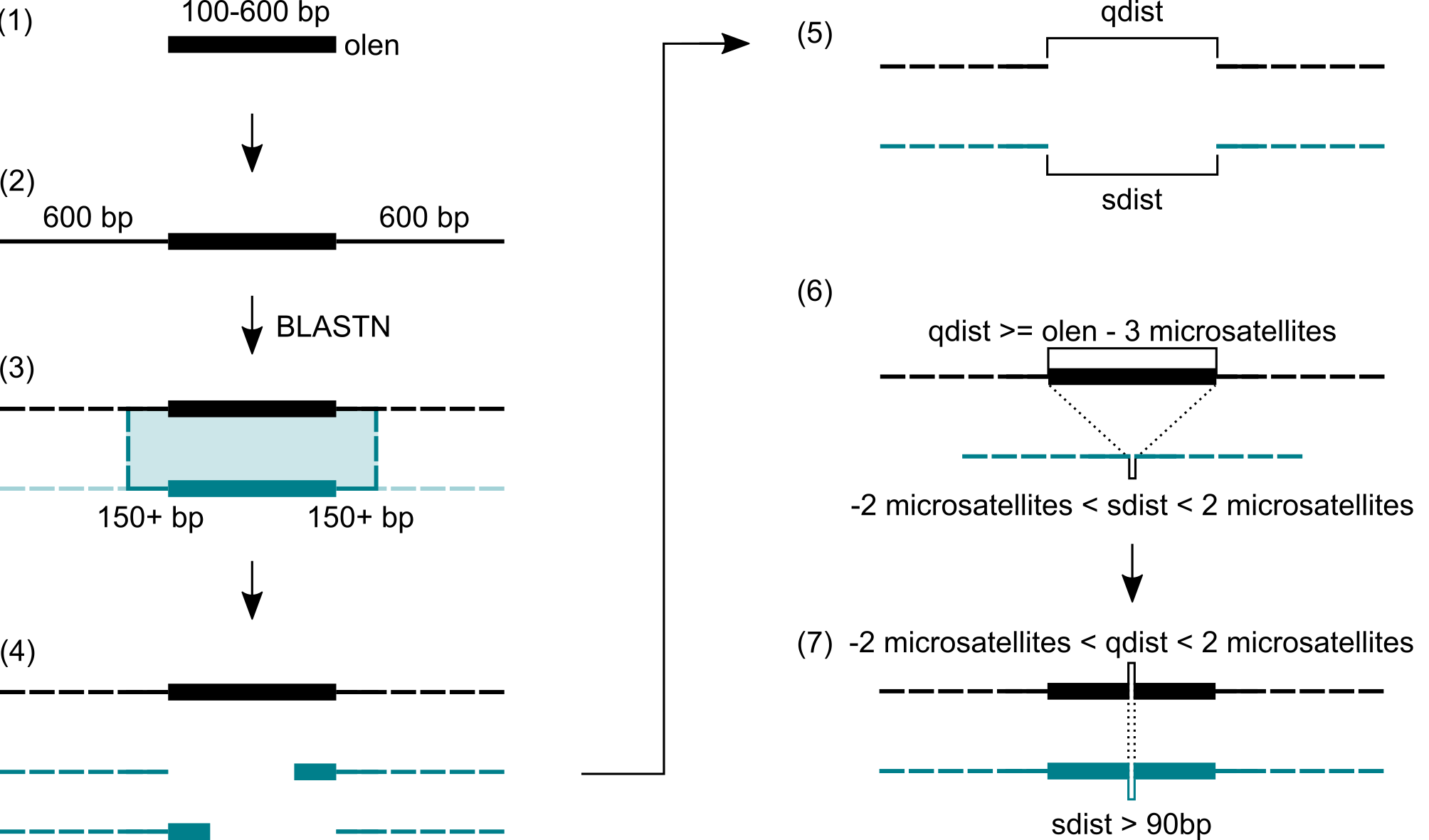

### SI_Fig_6_presence_absence_diagram.png

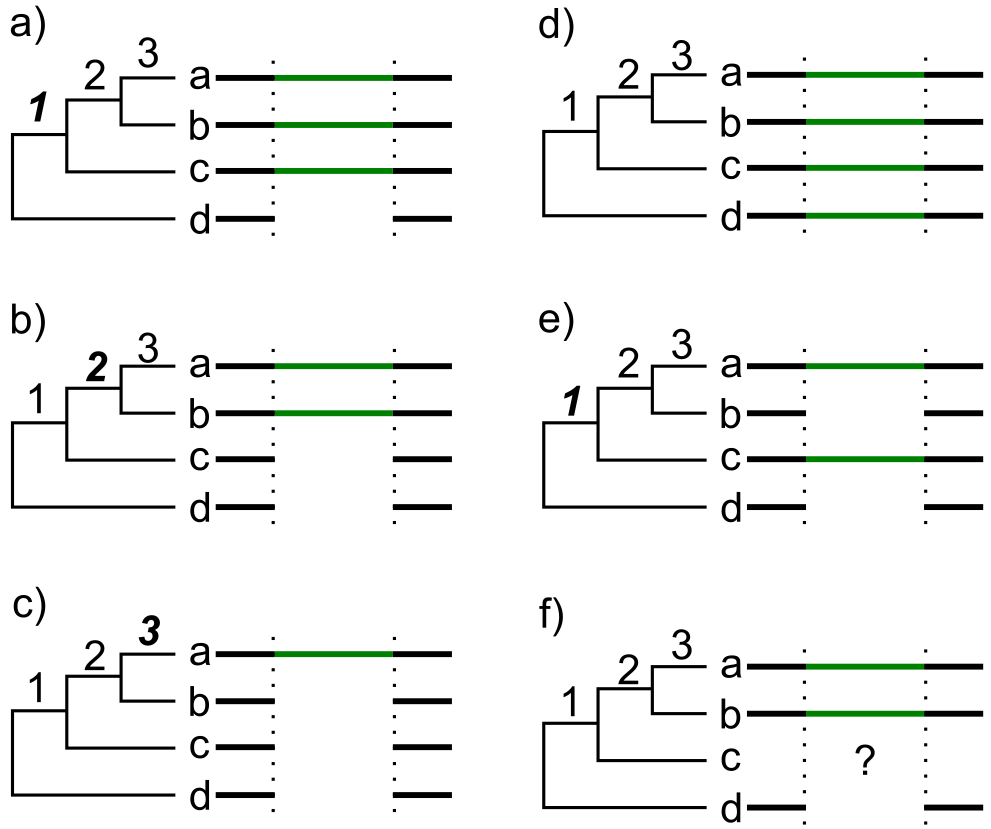
